# E-cadherin bridges cell polarity and spindle orientation to ensure prostate epithelial integrity and prevent carcinogenesis *in vivo*

**DOI:** 10.1101/245449

**Authors:** Xue Wang, Kai Zhang, Zhongzhong Ji, Chaping Cheng, Huifang Zhao, Yaru Sheng, Xiaoxia Li, Liancheng Fan, Baijun Dong, Wei Xue, Wei-Qiang Gao, Helen He Zhu

## Abstract

Cell polarity and correct mitotic spindle positioning are essential for the maintenance of a proper prostate epithelial architecture, and disruption of the two biological features occurs at early stages in prostate tumorigenesis. However, whether and how these two epithelial attributes are connected *in vivo* is largely unknown. We herein report that conditional genetic deletion of E-cadherin, a key component of adherens junctions, in a mouse model results in loss of prostate luminal cell polarity and randomization of spindle orientations. Critically, E-cadherin ablation causes prostatic hyperplasia which progresses to invasive adenocarcinoma. Mechanistically, E-cadherin and the spindle positioning determinant LGN interacts with the PDZ domain of cell polarity protein SCRIB and form a ternary protein complex to bridge cell polarity and cell division orientation. These findings provide a novel mechanism by which E-cadherin acts an anchor to maintain prostate epithelial integrity and to prevent carcinogenesis in vivo.

## Introduction

The prostate initially arises from embryonic urogenital sinus and undertakes ductal morphogenesis postnatally [1,2]. Murine prostatic epithelia are comprised of an inner single layer of polarized luminal cells, an outer layer of loosely distributed basal cells and a small fraction of scattered neuroendocrine cells [3,4]. Basal and luminal cells in the developing prostate epithelium display distinct cell division modes [5]. Luminal cells undergo symmetrical cell divisions during which the spindle orientation aligns parallel to the epithelial lumen and mother cell divides horizontally to generate two luminal cells. In contrast, basal cells undergo either horizontal symmetrical cell divisions to reproduce themselves or vertical asymmetrical cell divisions to give rise to a basal and a luminal daughter cell [5]. Horizontal cell division is of great importance for not only the surface expansion of prostate secreting lumen but also the maintenance of a monolayer luminal epithelial architecture, loss of which is an early event in prostate adenocarcinoma development. However, the molecular mechanism that ensures the horizontal symmetrical cell division of prostate luminal cells remains largely unknown.

Previous work has demonstrated that cell polarity is indispensible for correct cell division orientations. Cell polarity are instructed by three types of asymmetrically distributed polarity protein complexes, the Scribble (SCRIB)/Lethal giant larvae (LGL)/Discs large (DLG) protein complex beneath the basolateral cell membrane, the partitioning defective 3 (PAR3)/PAR6/atypical protein kinase (aPKC) in the cell apical-basal domain, and the Crumbs/PALS/PATJ protein complex under the apical cell membrane. Intensive studies in *Drosophila* have demonstrated that distribution cues for the spindle orientation determinants are derived from cell polarity [6]. An evolutionally conserved leucine-glycine-asparagine repeat protein (LGN)/nuclear and mitotic apparatus (NUMA)/ inhibitory alpha subunit of heterotrimeric G protein (Gαi) complex, which forms a lateral cortical belt to generate forces on spindle astral microtubules through interacting with dynein/dynactin, has been shown to be a major spindle positioning machinery [7]. Apical distribution of polarity protein aPKC phosphorylates LGN to exclude LGN from the apical cortex and determines the planar plane of the cell division [8–11]. DLG can interact directly with LGN and control its localization to orient the spindle position [12–14]. SCRIB or DLG knockdown in the developing *Drosophila* wing disc epithelium results in spindle orientation defects [15]. Nevertheless, how the polarity cues and horizontal spindle orientation are connected in mammalian systems is not fully understood.

Adherens junctions are closely associated with cell polarity and mitotic spindle positioning. They provide spatial landmarks for the anchorage of polarity proteins such as SCRIB and DLG [16]. Disrupted adherens junctions due to low calcium lead to diffused cell polarity protein localization in mammalian epithelial cells [17]. In neuroepithelial cells of *Drosophila*, adherens junction defects cause a shift of cell division modes due to deviation of spindle positioning from the planar axis and uncoupling of spindle positioning from asymmetric basal protein localization [18]. In the *Drosophila* follicular epithelium, adherens junctions prompt the apical position of the midbody to achieve asymmetric cytokinesis [19]. While these studies implicate the importance of adherens junctions in cell polarity and spindle positioning, the evidence for adherens junctions, such as E-cadherin, as a bridge between the two events is lacking.

In the present study, using a murine prostate-specific E-cadherin knockout model, we find that E-cadherin connects the cell polarity and spindle positioning to ensure the horizontal symmetric division of luminal cells and the integrity of prostate epithelia. Importantly, we show that a cell polarity protein SCRIB is recruited by E-cadherin to form an E-cadherin/SCRIB/LGN complex to guarantee the proper cell division mode and to prevent prostatic carcinogenesis.

## Results

### Loss of E-cadherin leads to a hyperproliferative phenotype of prostatic luminal cells and development of prostate adenocarcinoma in aged mice

To unravel the role of adherens junctions in prostate epithelial development and homeostasis, we first examined expression patterns of E-cadherin, a key component of adherens junctions, at different developmental stages via triple staining for E-cadherin, the basal prostatic cell-specific marker p63 and the luminal cell marker CK8. We found that E-cadherin was expressed throughout the epithelium at postnatal day 5 (P5), and then more predominantly expressed in the lateral cell membrane of tightly-contacted prostatic luminal cells than scattered basal cells over time (Supplementary Fig. 1A, B). Using the Cre-LoxP recombination system, we generated a mouse model in which E-cadherin was specifically knocked out in the prostate epithelium by crossing *Cdh1^fl/fl^* mice with probasin-cre transgenic mice. Immunostaining results confirmed an efficient E-cadherin deletion in the prostate of *Pcre;Cdh1^fl/fl^* mice (Fig. 1A, B). Through immunofluorescent staining of Ki67, a mitotic cell marker, we observed a significant increase of proliferative luminal cells from different lobes of *Pcre;Cdh1^fl/fl^* mice at P5, P10 and p15 or during adult prostate regeneration, compared with their littermate controls (Fig. 1C, D, Supplementary Fig. 2A, B and Supplementary Table 1). In contrast, significantly elevated cell proliferation was only detected in p63^+^ cells from P5 but not P10 or P15 *Pcre;Cdh1^fl/fl^* mouse prostates (Fig. 1C, D, Supplementary Fig. 2A, B, and Supplementary Table 1). This is possibly due to the minimal expression of E-cadherin in basal cells at P10 and P15. Those data suggest that E-cadherin deficiency leads to a hyperproliferative phenotype of prostatic luminal cells.

**Figure 1.**
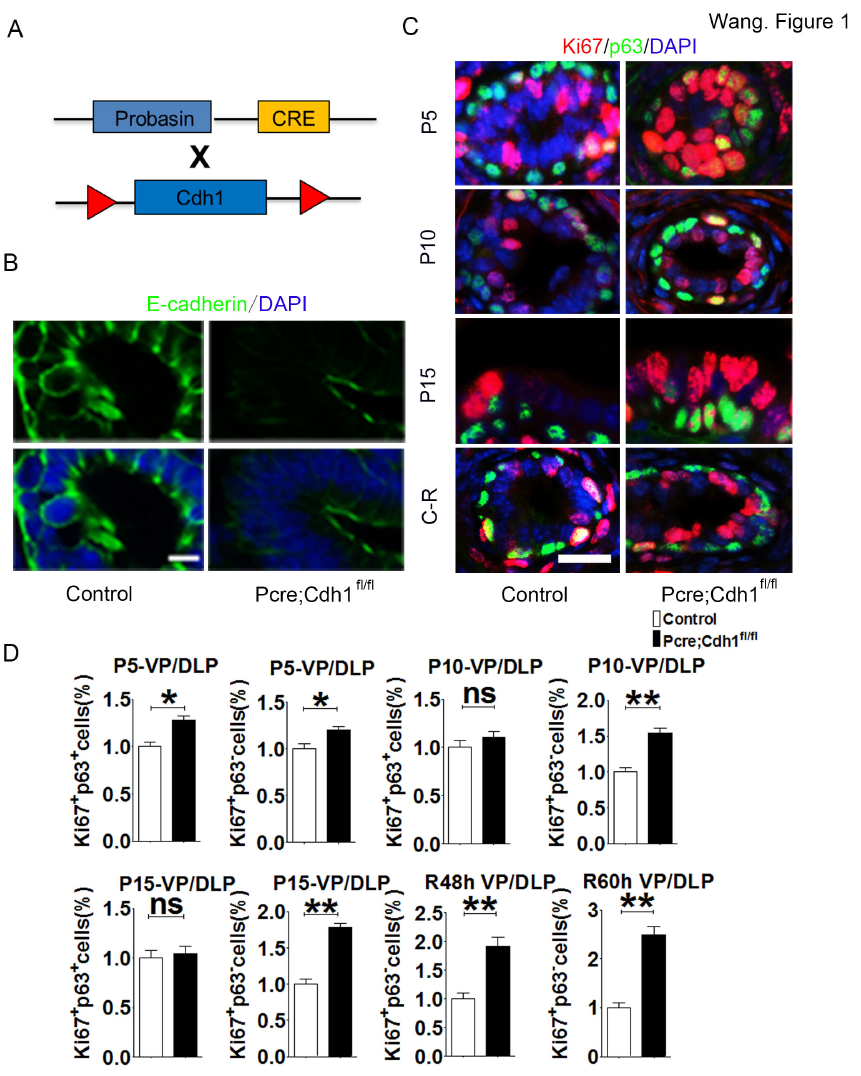
E-cadherin ablation leads to a hyperproliferative phenotype of prostatic luminal cells. (A)The schematic illustrates the generation of a mouse model with prostatic specific knockout of E-cadherin. (B)The efficiency of conditional E-cadherin deletion is confirmed by immunofluorescent staining. (C)Analysis of cell proliferation in developing prostates (P5, P10 and P15) or regenerating prostates (48h and 60h after androgen replacement) by immunofluorecent co-staining of proliferative marker Ki67 and basal cells marker p63 (Scale bars are 20μm for B and C). (D)Quantification of mitotic epithelial cells from ventral and dorsolateral prostatic lobes at different development stages or regeneration time points (n=3，Student’s *t*-test, ***P＜0.001,**P＜0.01, *P＜0.05,error bars = SEM.).

To determine whether this hyperproliferation led to prostate cancer initiation, we carefully followed prostates of *Pcre;Cdh1^fl/fl^* mice and their control littermates from 6 weeks to 21 months postnatally. Immunohistochemical staining analysis showed that at 6 weeks, different prostate lobes (anterior, ventral and dorsolateral lobes) of *Pcre;Cdh^fl/fl^*mice developed multifocal epithelial hyperplasia and multilayered polyp-like structure (Supplementary Fig. 2C). We detected low-grade murine intraepithelial neoplasia (mPIN) in 4 months old *Pcre;Cdh1^fl/fl^* mice (Fig. 2A). By 9 months of age, *Pcre;Cdh1^fl/fl^* mice developed high-grade mPIN, characterized by increased number of atypical cells with nuclear enlargement, prominent nucleoli and an elevated nuclear to cytoplasmic ratio (Fig. 2B). Of note, we detected invasive adenocarcinoma in the prostates of 21-month-old *Pcre;Cdh1^fl/fl^* mice manifested by loss of p63^+^ basal cells, disruption of the basement membrane, as demonstrated by loss of laminin staining and invasion of epithelial cells into the surrounding prostatic stroma (Fig. 2C-E). Further immunofluorecent staining confirmed the E-cadherin deletion in actively proliferating prostate epithelial cells from *Pcre;Cdh1^fl/fl^* mice. The vast majority of tumor cells in aged *Pcre;Cdh1^fl/fl^* prostates also displayed an absence of E-cadherin expression, indicating that the tumors were derived from hyperproliferation of E-cadherin knockout cells (Fig. 3A, B). However, E-cadherin deletion did not result in an epithelial to mesenchymal transition, as we did not observe an enhanced expression of mesenchymal cell marker vimentin in *Pcre;Cdh1^fl/fl^*prostate epithelial cells (Fig. 3C). Similar to human prostate cancer samples, we also occasionally detected apoptotic epithelial cells in *Pcre;Cdh1^fl/fl^* mouse prostate tumors (Fig. 3D).

**Figure 2.**
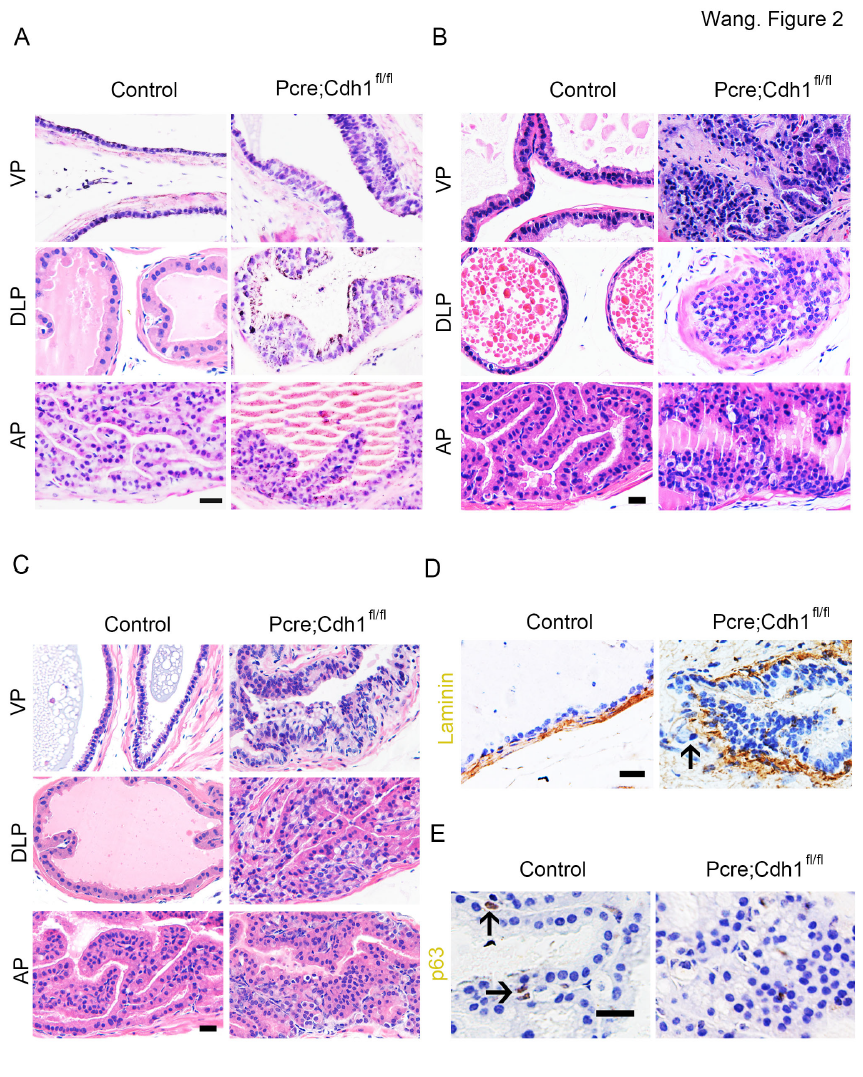
E-cadherin knockout results in development of invasive prostate adenocarcinoma. (A) H&E staining shows that multilayered epithelia structure and hyperplastic lesions can be found in 4-month old E-cadherin knockout mouse prostates. (B)Nine-month old E-cadherin knockout mouse prostates develop high-grade murine intraepithelial neoplasia. H&E staining shows adenocarcinoma in the prostates of 21-month-old *Pcre;Cdh1^fl/fl^*mice. (D)Immunohistochemistry staining of laminin indicates the disruption of the basement membrane and invasion of epithelial cells into the surrounding prostatic stroma in adenocarcinoma from *h1^fl/fl^* prostates. (E)Loss of p63^+^ basal cells in adenocarcinoma from *Pcre;Cdh1^fl/fl^* prostates (Scale bars are 20μm).

**Figure 3.**
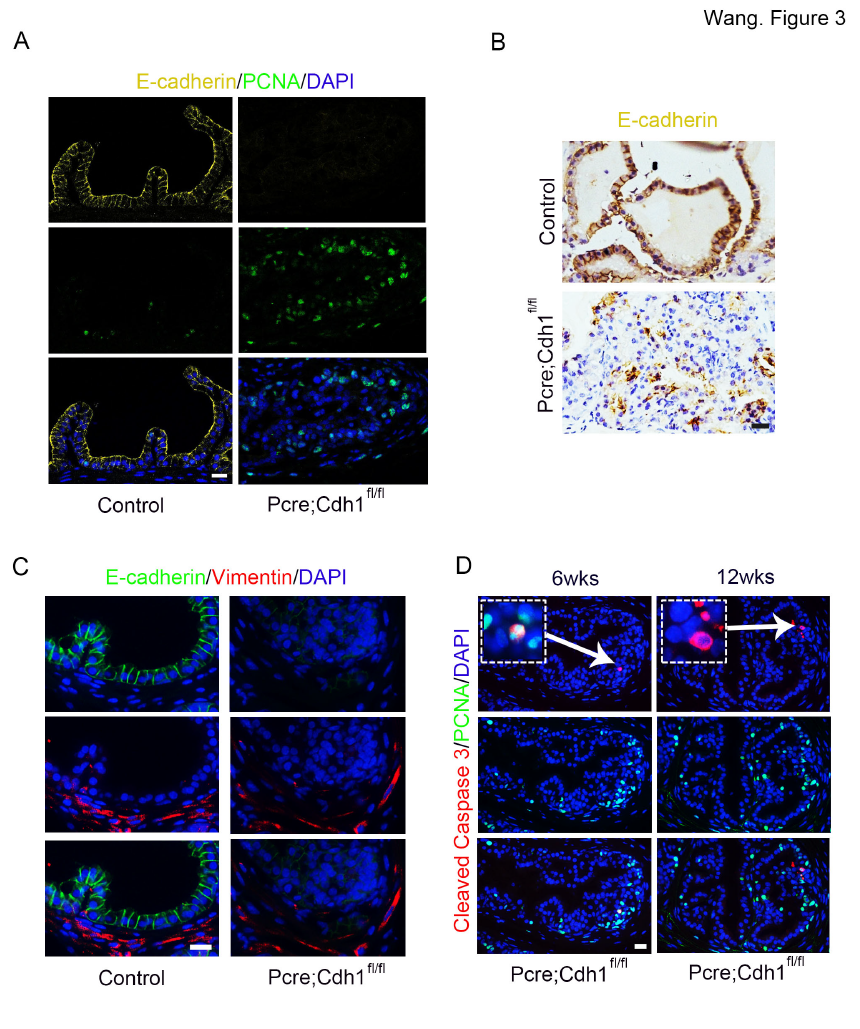
Tumors in *Pcre;Cdh1^fl/fl^* prostates are derived from hyperproliferation of E-cadherin knockout cells. (A)IF staining confirms E-cadherin deletion in actively proliferating prostate epithelial cells of E-cadherin knockout mice. (B)Tumor cells in aged E-cadherin knockout prostates display absence of E-cadherin expression. (C)E-cadherin ablation does not result in an epithelial to mesenchymal transition (EMT). (D)Apoptotic epithelial cells are occasionally detected in hyperproliferative multilayered regions of E-cadherin knockout mouse prostates (Scale bars are 20μm).

### Horizontal division of prostate luminal cells is randomized by E-cadherin deletion during postnatal development and regeneration

Previous studies have shown that planar spindle alignment in epithelia is crucial for the maintenance of mitotic daughter cells within the plane of the tissue, whereas spindle orientation perpendicular to the basement membrane is required for asymmetric cell divisions and epithelial stratification in tissues such as skin [20]. The expansion of a monolayer developing prostate lumen is ensured by the horizontal division of luminal cells [5], disruption of which often causes multilayered polyps-like structure as detected in *Pcre;Cdh1^fl/fl^* prostates. We therefore wondered whether loss of E-cadherin affected proper spindle alignment of dividing luminal cells during postnatal prostate development. To test this possibility, we analyzed the spindle orientation of dividing luminal cells by staining sections of developing prostates with p63 and survivin, an approach used for analysis of cell division modes [5]. We observed a more than 2-fold increase of tilted and vertical cell divisions in luminal cells in P5 E-cadherin knockout prostates (38.46% in *Pcre;Cdh1
^fl/fl^* mice versus 14.85% in control mice). Similar findings were obtained from P10 and P15 prostates (Fig. 4A, C, D and Supplementary Table 2). In addition, during adult prostate regeneration, E-cadherin deletion resulted in randomization of the luminal cell division plane (Fig. 4B, E and Supplementary Table 2). Collectively, these findings suggest that E-cadherin is required for a proper mitotic spindle orientation during prostate development and regeneration.

**Figure 4.**
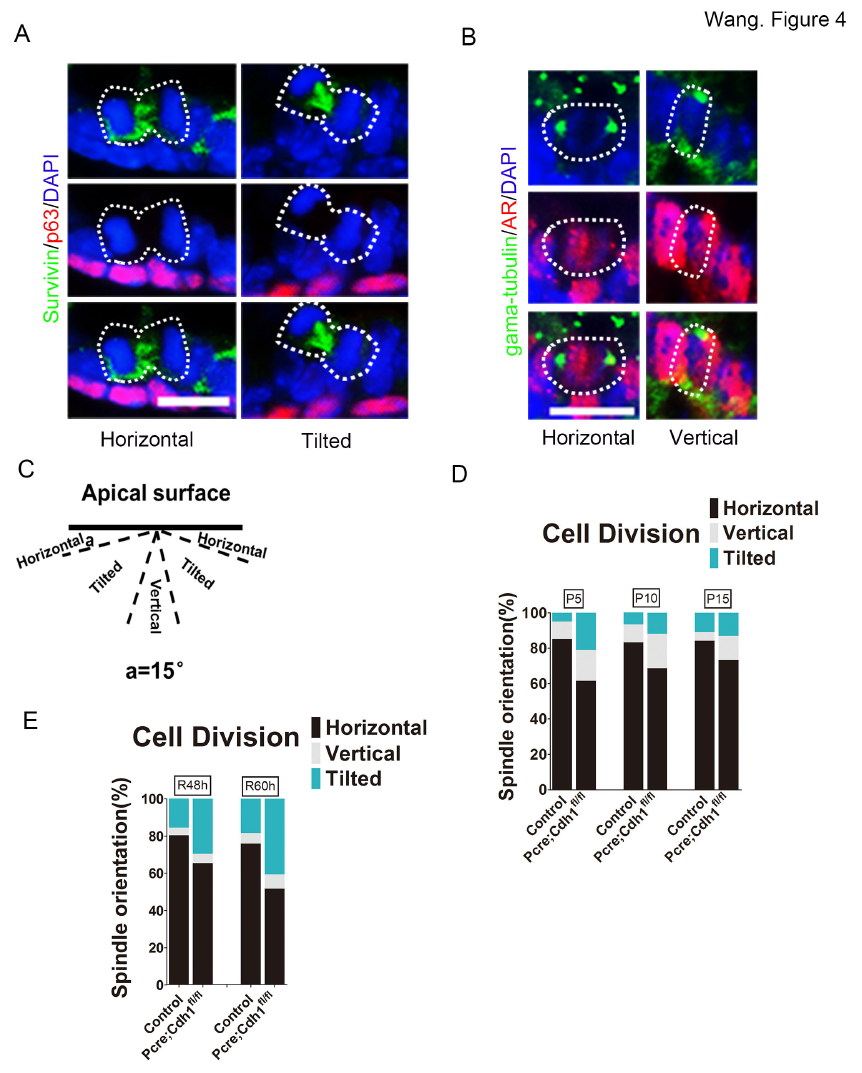
Oriented division of prostate epithelial cells during postnatal development and regeneration is randomized by E-cadherin deletion *in vivo*. (A)Immunofluorecent staining of p63, survivin and DAPI illustrates mitotic spindle orientation of dividing prostate epithelial cells. (B)Co-staining of AR, gama-tubulin and DAPI shows that prostate luminal cells mostly undergo horizontal divisions and very rarely go through vertical divisions during prostate development (Scale bars are10μm for A and B). (C)The schematic in the left panel depicts the definition of horizontal, tilted or vertical spindle orientation based on the angles of spindle alignment relative to the apical surface. Division planes positioned at 75–90 degrees relative to the basement membrane were defined as vertical divisions, those that were oriented at 15–75 degrees were classified as tilted divisions, while those that were oriented at 0–15 degrees were considered as horizontal divisions. (D-E) E-cadherin deletion leads to a significant increase in vertically or obliquely dividing luminal cells during prostate development (D) or regeneration (E).

### Interference of E-cadherin expression results in spindle dis-orientation in the prostatic RWPE-1 cell line

To directly test whether E-cadherin loss led to defects in spindle positioning and subsequently cell division orientation, we modified a previously reported approach to record the spindle position of an immortalized prostate epithelial cell line RWPE-1 in real time [21,22]. RWPE-1 cells were co-transfected with an H2B-RFP and an α-tubulin-GFP plasmid to mark chromosomes and spindles respectively (Fig. 5A). Knockdown of E-cadherin was achieved by infection of RWPE-1 cells with a shRNA containing lentiviral construct (Fig. 5B). Control or E-cadherin knockdown cells were plated on retronectin-coated dishes and imaged under a confocal microscopy equipped with a live-cell recording unit (Fig. 5C). Four-dimensional movies of dividing RWPE-1 cells were analyzed to measure angles of mitotic spindles relative to the culture dish. In line with previous reports [23,24], we found that mitotic spindles of control RWPE-1 cells were positioned at a wide range of angles during prometaphase, but the majority of spindles (83.5%) exhibited a parallel orientation to the substrate from the late metaphase to telophase (Fig. 5D, E, Supplementary Table 2 and Supplementary Movie 1). This horizontal position of spindles ensured a cell division cleavage plane perpendicular to the substrate. However, we detected a striking increased percentage of E-cadherin-knockdown cells which could not correctly position their spindles in telophase, thereby caused a randomized cell division cleavage plane (Figure 5D, E, Supplementary Table 2 and Supplementary Movie 1). These *in vitro* experiments substantiated our *in vivo* observations that adherens junction protein E-cadherin was indispensable for the proper positioning of mitotic spindles in prostate epithelial cells.

**Figure 5.**
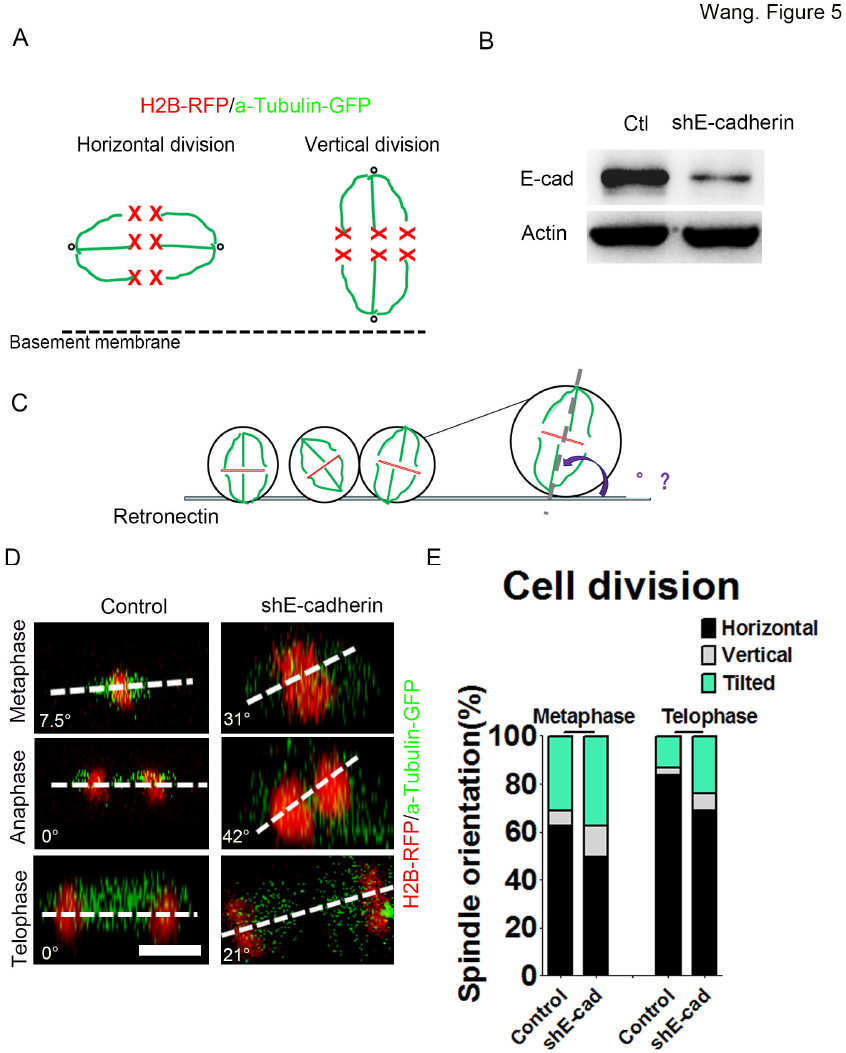
E-cadherin knockdown causes mitotic spindle disorientation in the immortalized prostate cell line RWPE-1. (A)The cartoon illustrates different division modes of epithelial cells. The green lines are α-tubulin-GFP marked spindles and the red crosses depict H2B-RFP labeled chromosomes. (B)The efficiency of E-cadherin knock-down is confirmed by Western blotting. (C)The cartoon illustrates how the mitotic spindle angle of cultured RWPE-1 cells relative to the retronectin-coated dish bottom is measured. (D)Representative confocal images of α-tubulin-GFP marked spindles and H2B->RFP labeled chromosomes from control or E-cadherin knockdown RWPE-1 cells in different cell cycle phases (Scale bars are 20μm). (E)Quantification of metaphase and telophase spindle angles indicated that E-cadherin knockdown causes a remarkable increase of RWPE-1cells with vertical or oblique mitotic spindle alignment in the telophase.

### Ablation of E-cadherin disrupts cellular distribution and association of key spindle positioning proteins LGN and NUMA

Luminal epithelial cells of a developing prostate divide within the plane of the epithelium by orienting spindle poles towards the lateral membrane. Intensive research has revealed that an evolutionarily conserved LGN/NUMA complex plays an essential role in directing the cortical attachment of spindle astral microtubules to dynein in variety tissues from both invertebrates and vertebrates [9,24–26]. In agreement with previous reports, we found that LGN and NUMA formed two crescents underneath the lateral membrane and in parallel to the basal surface in mitotic wild-type prostate luminal cells (Fig. 6A-D). However, in the absence of E-cadherin, distribution of LGN and NUMA in dividing luminal cells can be detected all around the cell cortex (Fig. 6A-D and Supplementary Table 3), suggesting that disoriented mitotic spindle alignment was due to the diffused localization of LGN/NUMA complex. In addition, the protein-protein interaction between LGN and NUMA was also severely abrogated after E-cadherin deletion (Fig. 6E). As a result, the luminal cells underwent perpendicular or oblique divisions, which disrupted the monolayer epithelial structure and formed the disorganized, multilayered polyps-like structure (Fig. 1 and Supplementary Fig. 2).

**Figure 6.**
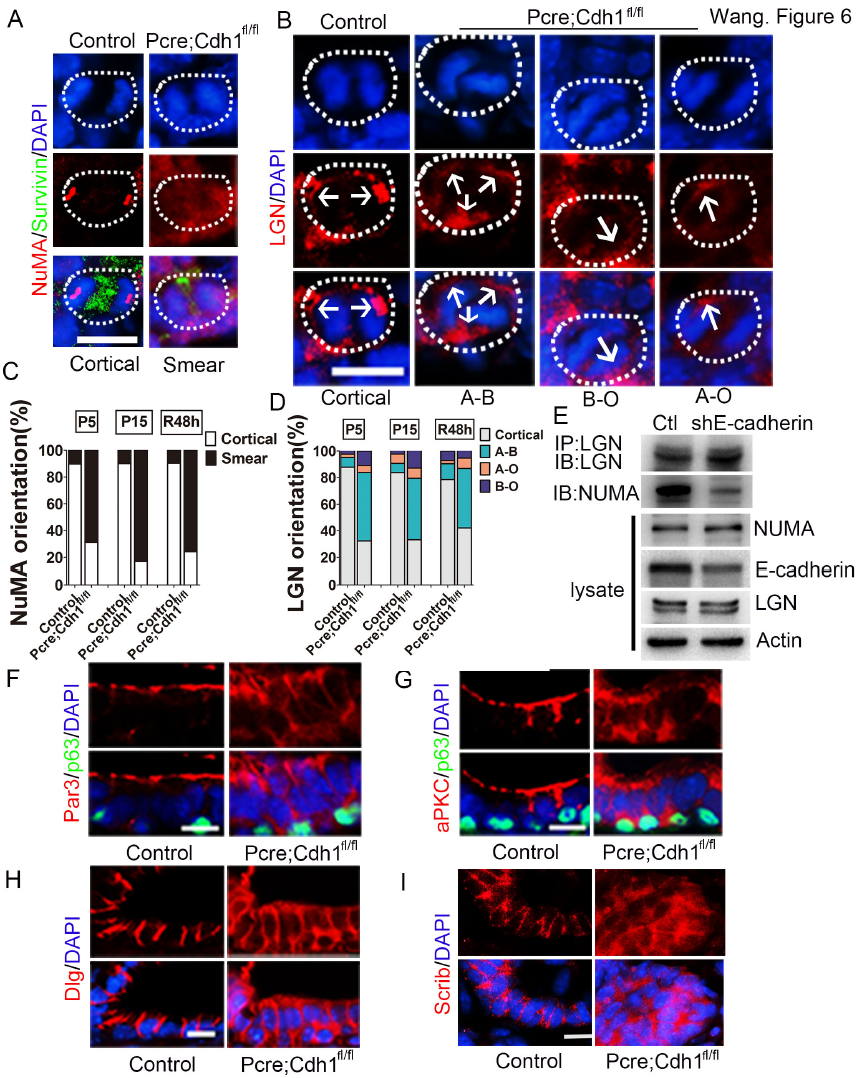
Ablation of E-cadherin disrupts the distribution of key spindle positioning and cell polarity proteins of prostate luminal cells. (A)Immunofluorecent staining of NUMA shows that E-cadherin depletion in the mouse prostate diminishes NUMA cortical enrichment. (B)LGN can be detected all around the cell cortex of dividing E-cadherin knockout prostate epithelial cells, in comparison to its exclusively lateral distribution in wild-type controls. A-B, diffused LGN position; B-O, basal surface enriched LGN position; A-O, apical surface enriched LGN position. (C)Quantification of the NUMA localization pattern in dividing luminal cells during prostate development and regeneration. (D)Quantification of ratios of each type of LGN distributions in dividing luminal cells during prostate development and regeneration. (E)The formation of LGN/NUMA protein complex is markedly suppressed due to E-cadherin knockdown *in vitro*. (F-G)Wild-type prostate luminal cells exhibit apical distribution of polarity proteins PAR3 (F) and aPKC (G), whereas E-cadherin deleted luminal cells display a diffused expression pattern of those polarity proteins. (H-I) Immunostaining of basolateral polarity proteins, DLG (H) and SCRIB (I) shows that the basolateral polarity of luminal cells is dispersed by E-cadherin ablation. All sections were collected from P15 prostate tissues and counterstained by 4’,6-diamidino-2-phenylindole(DAPI)(blue) (Scale bars are10μm).

### Loss of E-cadherin impairs the cell polarity of prostate luminal cells during prostatic development

Cell polarity provides spatial cues to guide spindle orientation [10,12,14,27]. Disruption of cell polarity is often seen during the course of cancer initiation [11,28–32]. Given the observation that horizontal mitotic spindle positioning of prostate luminal cells was severely disrupted in the *Pcre;Cdh1^fl/fl^* prostate, we therefore wondered whether E-cadherin affected luminal cell polarity. We next investigated the impact of E-cadherin loss on the expression and distribution of PAR complex and SCRIB complex. Utilizing immunofluorecent staining, we detected core components of the PAR complex, PAR3 and aPKC were localized in the apical domain of wild-type prostate luminal cells, but diffusedly distributed in *Pcre;Cdh1^fl/fl^* prostate luminal cells (Fig. 6F, G and Supplementary Fig. 3A, B). Likewise, the normal basolateral distribution of polarity proteins DLG-1 and SCRIB in prostate luminal cells were markedly impaired after E-cadherin deletion (Fig. 6H, I and Supplementary Fig. 3C, D). In addition, the formation of SCRIB/DLG polarity protein complex was disrupted following E-cadherin deletion (Supplementary Fig. 3E). Together, these results indicated that loss of E-cadherin exerted a devastating influence on the formation and maintenance of prostate luminal cell polarity.

### E-cadherin recruits SCRIB to form an E-cadherin/SCRIB/LGN complex to bridge cell polarity and spindle positioning

Previous research has demonstrated that adherens junctions are necessary for the establishment of cell polarity and positioning mitotic spindles symmetrically in *Drosophila* [18]. Given the aforementioned findings that E-cadherin deletion or knockdown led to cell polarity loss and spindle dis-orientation, we proposed a hypothesis that E-cadherin may serve as a central molecule to bridge cell polarity and spindle positioning. It was reported that in mammalian epithelial cells, restriction of SCRIB to the lateral cell-cell junction is dependent on E-cadherin and that SCRIB reciprocally regulates E-cadherin–mediated cell adhesion [16,33,34]. On the other hand, lateral localization of the SCRIB polarity complex determines the planar orientation of mitotic spindles [14,15,35,36]. Therefore, we then tested whether there were physical interactions between E-cadherin and SCRIB. Co-IP assay showed endogenous interactions between E-cadherin and SCRIB in RWPE-1 cells (Fig. 7A, B). In addition, we observed that E-cadherin associated with LGN (Fig. 7A, C). To elucidate whether possible interactions existed between the SCRIB complex and the LGN complex in prostate epithelial cells, we carried out additional co-IP experiments. Intriguingly, we found that endogenous SCRIB interacted with LGN, which can be suppressed by E-cadherin knockdown (Fig. 7B-D). Furthermore, the association between LGN and E-cadherin was markedly attenuated by SCRIB knockdown (Fig. 7E). To determine whether the interactions between E-cadherin and SCRIB, as well as LGN with SCRIB were direct, and which domains of SCRIB were responsible for the interactions, we designed and performed GST-pull down experiments with different truncated fragments of SCRIB. As show in Fig. 7F-G, the fragment containing the PDZ domain of SCRIB directly bound to both the E-cadherin and LGN. Collectively, these data revealed a ternary protein complex of E-cadherin/SCRIB/LGN by which adherens junctions were able to tightly regulate and efficiently connect cell polarity complexes and spindle orientation determinants so to precisely position mitotic spindles in dividing cells (Fig. 8).

**Figure 7.**
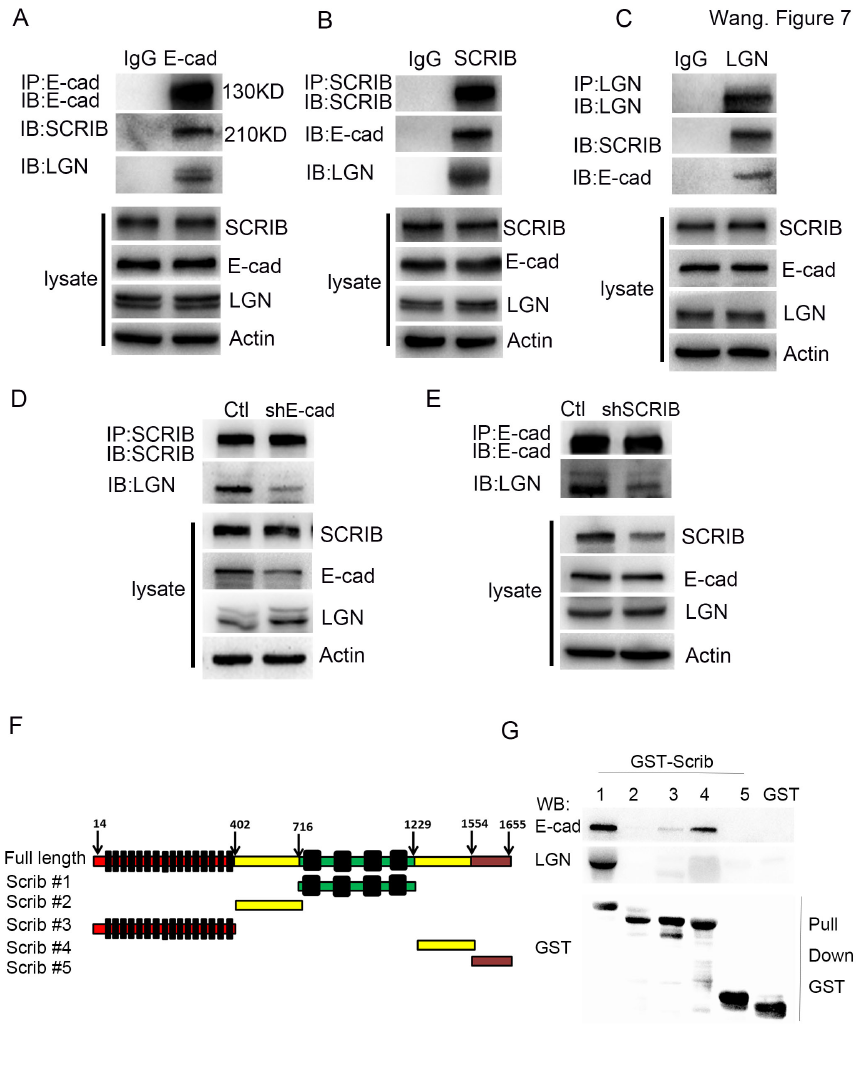
E-cadherin recruits SCRIB to form an E-cadherin/SCRIB/LGN complex to connect cell polarity and spindle positioning. (A)Co-IP experiments using RWPE-1 cell lysates suggest that endogenous E-cadherin interacts with SCRIB and LGN. (B)Endogenous SCRIB co-immunoprecipitates with the E-cadherin/LGN protein complex. (C)Co-IP assays indicate that endogenous SCRIB, E-cadherin and LGN form a ternary complex. (D)The interaction between SCRIB and LGN is suppressed by E-cadherin knockdown. (E)SCRIB knockdown attenuates the association between LGN and E-cadherin. (F)Diagram of SCRIB fragments used to identify E-cadherin and LGN binding domains. (G)Pull down assays indicate the fragment containing PDZ domain of SCRIB binds to E-cadherin and LGN.

**Figure 8.**
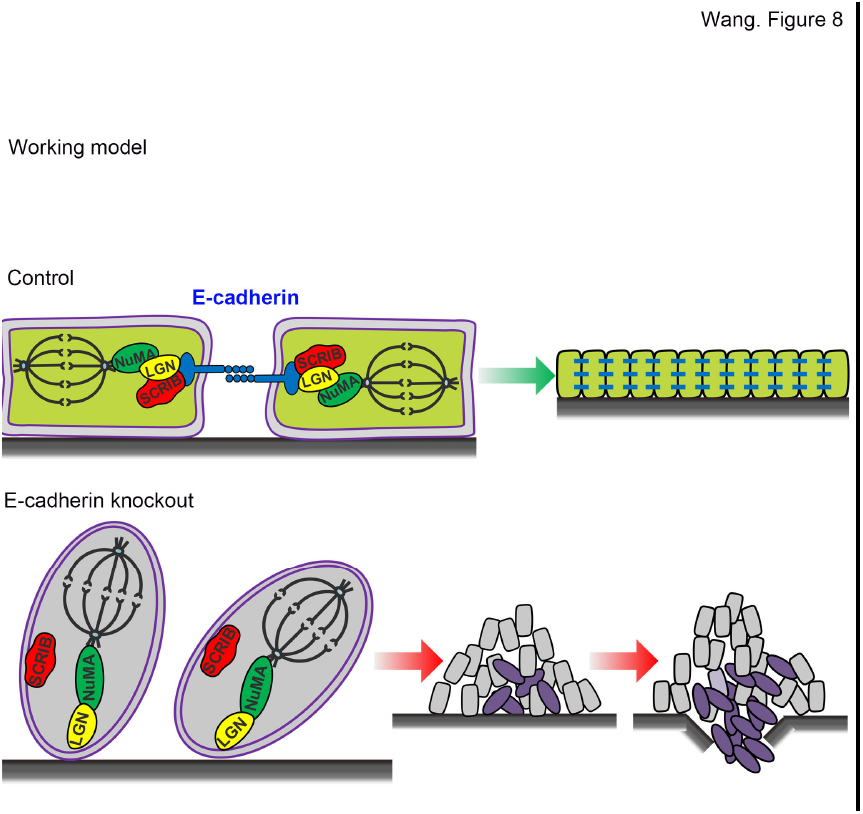
The working model shows adherens junction protein E-cadherin recruits SCRIB to the LGN/NUMA complex to connect the cell polarity ad mitotic spindle positioning. E-cadherin loss leads to disruption of prostate luminal cell polarity and randomization of spindle orientations, which strongly predispose mice for prostate tumorigenesis.

## Discussion

At the current study, we find that genetic deletion of E-cadherin in the mouse prostate results in cell polarity loss, mitotic spindle dis-orientation, and monolayer structure disruption in the luminal epithelium. Importantly, E-cadherin knockout leads to prostatic hyperplasia which progresses to invasive adenocarcinoma in aged mice. Mechanistically, E-cadherin can recruit SCRIB to form a protein complex with LGN by directly binding to the PDZ domain of SCRIB to restrict SCRIB and LGN to the lateral cell membrane. These findings provide a novel mechanism that E-cadherin acts as a vital bridge between cell polarity and spindle orientation for the maintenance of a normal prostate epithelial architecture.

In both *Drosophila* and mammalian systems, adherens junctions, cell polarity and mitotic spindles orientation are intricately intertwined biological processes. However, how these three fundamental attributes of the epithelium are connected remains elusive. In particular, limited previous studies regarding this question were mostly carried out in cell lines or in invertebrates. The current study was carried out in a genetically engineered mouse model. Our examination of early murine postnatal prostate development and adult regeneration reveals that E-cadherin deletion results in loss of luminal cell polarity and mitotic spindle dis-orientation *in vivo*. In addition, other than microscopic examination of cell mitosis at limited time points, we used a real-time recording system which allows us to follow the whole process of individual cell division. By this way, we further demonstrate vividly that E-cadherin is required for the horizontal alignment of spindles in the telophase of dividing RWPE1 cells *in vitro*. Critically, we uncover a central role of adherens junction protein E-cadherin in the coordination of cell polarity and mitotic spindles orientation by forming an E-cadherin/SCRIB/LGN protein complex. E-cadherin knockout leads to a dis-assembly of the SCRIB polarity complex and the LGN/NUMA complex, causing a subsequent randomization of the luminal cell division plane and forming of multilayered, disorganized epithelia, a common feature of the early stage of prostate tumorigenesis.

Based on biochemical analysis and examination of MDCK and U2OS cell lines, Nelson, W.J.’s group reported recently that E-cadherin instructs cell division orientation by binding to LGN and brings LGN to cell-cell adhesions [37]. In a related study on MDCK cells, Hart, K.C. et al reported that E-cadherin acts to sense mechanical tension across an epithelial sheet and facilitate polarized cortical distribution of LGN to align cell divisions[38]. However, whether this LGN/E-cadherin complex is important for epithelial tissue integrity and the consequences of disruption of this interaction *in vivo* is unknown. In addition, whether additional components, such as cell polarity, are required to the intricate regulation of LGN by E-cadherin is not determined. Our current study shed new important lights on these aspects:1) We demonstrated clearly *in vivo* that E-cadherin knockout causes LGN/NUMA dissociation and dis-localization during luminal cell mitosis, which subsequently leads to luminal cell division plane dis-orientation and serious impairment of normal prostate architecture; 2) We provide direct evidences that cell polarity protein SCRIB is required for the efficient complex-forming between E-cadherin and LGN in prostate epithelial cells, as SCRIB knockdown markedly reduces the association of E-cadherin and LGN. Moreover, we found that both E-cadherin and LGN can directly bind to the fragment containing PDZ domain of SCRIB by pull down assays. Of note, our experiments also provide a novel molecular explanation for previous reports regarding how SCRIB is required for the establishment of E-cadherin–mediated cell–cell adhesion and correct positioning of mitotic spindles [16,33,34].

Defects in precisely controlled cell-cell adhesion and junctions, cell polarity and mitotic spindles position are closely associated with developmental disorders, and may contribute to cancer initiation and progression [31],39,40]. For example, decreased expression or genetic loss of adherens junction molecule E-cadherin is frequently found in various cancer types including gastric carcinomas, lung cancer and breast cancer [41–43]. In prostate cancer, E-cadherin down regulation is significantly associated with advanced stages and tumor metastasis [44,45]. A recent study using a NKX3.1 CRE-ERT2;Cdh1^fl/fl^ mouse model showed that tamoxifen induced partial E-cadherin deletion in luminal cells from 8-week old mice leads to a short-term of anoikis [46]. While we do also detect some apoptotic cells in hyperplastic lumen of *Pcre;Cdh1^fl/fl
^* prostates in this study, we demonstrate that a more efficient prostate epithelium deletion of E-cadherin leads to luminal epithelial cell hyperplasia. Those apoptotic luminal cells may come from actively perpendicular or oblique dividing cells that lose cell-cell contact or cell-basement membrane contact. Apoptotic luminal cells can also be frequently found in human prostate cancer samples. A similar simultaneous increase in both proliferation and apoptosis was reported in PAR3 deletion-induced mammary gland hyperplasia [32]. In addition, our observation that invasive prostate carcinoma develops in old E-cadherin knockout mice is in line with a clinical correlation between E-cadherin deletion or downregulation and human prostate cancer progression [44,45]. Moreover, immunofluorecent staining of developmental prostate or tumor tissues from *Pcre;Cdh1^fl/fl^* mice confirms E-cadherin deletion in most proliferating luminal cells, suggesting that the hyperplasia is not derived from compensatory proliferation of residue E-cadherin expressing cells. Therefore, loss of E-cadherin induced cell polarity and cell division plane deregulation can strongly predispose mice for prostate tumorigenesis. These findings provide evidence for an important role of E-Cadherin not only in anchorage of cell polarity proteins with spindle positioning determinants, but also in prostatic carcinogenesis.

## Methods

### Experimental Animals

*Probasin-Cre* and *Cdh^fl/fl^* mice were introduced from National Cancer Institute (NCI:01XF5) and Jackson laboratory (JAX:005319) respectively. All mice were maintained and utilized according to the ethical regulations at Ren Ji Hospital. The animal protocols were approved by the Ren Ji Hospital Laboratory Animal Use and Care Committee. Prostates of male mice were dissected for histology and immunostaining.

### Human Cell Lines

Prostate immortalized epithelial cell line RWPE-1 cells (CRL-11609) were cultured in defined keratinocyte-SFM basic medium with growth supplements (Giboco, 10744-019). Cells were propagated in an incubator with 5% CO2 at 37℃.

### Plasmids and Lentivirus production

The shRNA against E-cadherin, SCRIB and scramble shRNA were cloned into a lentiviral vector pLVTH. Lentivirus was produced as previously reported[47]. RWPE-1 cells were infected with lentivirus and selected with 3ug/ml puromycin for 2weeks to generate the stable E-cadherin and SCRIB knockdown cell line, respectively. The H2B-RFP (#26001) and a-tubulin-GFP (#64060) plasmids were purchased from Addgene, which were originally deposited by Dr. Beronja S. and Dr. Yang W..

### Castration and androgen replacement

*Pcre;Cdh1^fl/fl^* mice and their control littermates were surgically castrated. Three weeks after castration, dihydrotestosterone (MCE, HY-A0120) dissolved in sterile corn oil was given via intraperitoneal injection twice each day (50ug/d) to induce prostate regeneration. Prostate tissues were dissected for section and immunofluorecent staining at different regeneration stages (48 hours or 60 hours post testosterone administration).

### Immunofluorescence staining

Mouse prostates were fixed in 4% paraformaldehyde for 20 minutes and dehydrated in 30% sucrose solution overnight. Tissues were then embedded in Optimal Cutting Temperature (O.C.T.) compound and quickly frozen in a -80℃ refrigerator for 10min. Frozen sections were cut at a thickness of 6μm. The sections were washed with PBS and subjected to a heat-induced epitope retrieval step in 0.01M sodium citrate (PH 6.0). Then sections were transferred to a blocking solution (PBS with 0.2% TritonX-100 and 10% donkey serum) for 1 hour at room temperature. Primary antibodies diluted in PBS with 0.2% TritonX-100 and 1% donkey serum were then applied to the section overnight at 4℃ and washed away with PBS three times. Sections were then incubated with secondary antibodies, conjugated to Alexafluo-488, 594 or 546 for 1 hour at room temperature. After thorough washing, sections were mounted with Vector Shield mounting medium containing DAPI.

### Immunohistochemical staining

Paraffin-embedded tissue sections were deparaffinized, rehydrated and subjected to a heat-induced epitope retrieval step in 0.01M sodium citrate (PH 6.0). Endogenous peroxidase activity was ablated by using 3% hydrogen peroxide. Sections were blocked, stained with primary antibodies then horseradish peroxidase conjugated secondary antibodies as described above. Then DAB staining was undertaken according to the manufacturer’s instructions (DAB Staining kit, GK347010, Gene Tech (Shanghai) Company Limited). Sections were washed under running tapping water for 5min then counterstained with hematoxylin, followed by dehydration and mounting with the neutral balsam mounting medium.

### GST-Pull down

Human SCRIB full length was cloned from the cDNA of the RWPE-1 cell line. GST-tagged fragments of human SCRIB were cloned into the pET-49b(+)vector with SpeI and EcoRI restriction enzymes. DNA constructs were transformed into BL21 (DE3) E.coli for following recombinant protein expression. For protein purification, glutathione sepharose beads (GE Healthcare) were added to the recombinant protein containing bacterial lysate and incubated for 1.5 hours at 4℃. Beads were washed with 500µl cold bacteria lysis buffer twice. RWPE-1 cell lysate was added to the beads and incubated overnight at 4℃. After thorough wash, protein loading buffer were added to the beads and boiled for 10 minutes at a 100℃ metal bath. The immunoprecipitates were analyzed by SDS-PAGE and immunoblotting with GST, or indicated antibodies.

### Immunoprecipitation and Immunoblotting

For immunoprecipitation, RWPE-1 cells were washed with ice-cold phosphate-buffer saline and lysed in a lysis buffer (50mM Tris-HCL,150mM NaCl, 1mM EDTA, 0.25% TritonX-100, PH7.4) supplemented with protease and phosphatase inhibitors (Roche, Penzberg, Germany). Cell lysates were incubated with 1μg primary antibodies for 6 hours at 4℃.Rabbit or mouse immunoglobulin G(Sigma-Alrich, USA) was utilized as controls. Activated Protein G-Agarose (Roche, 11243233001) were added into the protein lysate and incubated for another 1 hour at 4℃. Then the immunoprecipitates were washed 3 times with the lysis buffer and denaturalized for 10 minutes at a 100℃ metal bath. The Immunoprecipitation results were detected by immunoblotting assay. Protein samples were separated via SDS-PAGE electrophoresis and then transferred to PVDF membranes. The membranes were blocked with 5% non-fat milk in TBST (TBS containing 0.1% Tween 20) and incubated with primary antibodies at 4℃ overnight. Membranes were then rinsed with TBST for 3 times and incubated with secondary antibodies for 1 hour at room temperature. After washing with TBST 3 times, membranes were exposed with enhanced chemiluminescence substrates (Thermo Scientific). Antibodies used in the study were listed in Supplemental Table S4.

### Confocal microscopy for images and videos

Lentivirus-transfected RWPE-1 cells were seeded on glass-bottom dishes coated with 0.1μg/μl Retronectin (Takara Bio). Images were acquired every 6-7 minute with a xyzt acquisition mode using an Axio Observer under a Z1 microscope with the LSM 700 scanning module (Zeiss). Cell cultures were maintained at 37℃ and 5% CO2 incubator for live imaging.

### RNA extraction and quantitative-PCR analysis

To collect single prostate epithelial cells suspension for flow cytometric sorting of basal, luminal and stromal cells, prostates from 12-week-old Pcre;Cdh1fl/fl mice and control littermates were harvested, minced and incubated with 3ml 1mg/ml Collagenase solution on a shaker at 37℃for 2 hours. The cells were then spin down and further digested with 2ml 0.25%Trypsin/EDTA at a 37℃ water-bath for another 6minutes. Single cells were obtained by filtering the mixture through a 40μm cell strainer. Percp-lineage, FITC-CD49f, APC-Sca-1 antibodies were applied to the cell suspension and incubated in the dark for 40 minutes. Basal (Lineage^-^Sca-1^+^CD49f^+^), luminal epithelial cells (Lineage^-^Sca-1^-^CD49f^+^)andstromalcells (Lineage^-^Sca-1^+^CD49f^-^)were sorted using a BD FACSARIA system. Total RNA were extracted from each cell population using Trizol reagents following instructions from the manufacturer (Life technologies).cDNA was synthesized from the extracted total RNA using PrimeScript RT Reagent Kit(RR037A). qPCR was performed using SYBR Premix Ex Taq (TliRNaseH Plus) (RR420A).Relative transcript abundance was determined by the comparative CT method using Actin as a reference gene. Primers used in RT-PCR were listed in Supplemental Table S5.

### Quantification and statistical analysis

The ImageJ 1.46r software was used to determine the positive stained cells in immunostaining images and measure angles of mitotic spindles relative to the retronectin base. Microsoft Excel and Graph Pad Prism5 were used for data compilation and graphical representation. All bar graphs, line graphs and dot plots are represented as mean ± standard deviation. All statistical analysis was done using a two-tailed Student’s *T-test* and a p-value<0.05 was considered significant and indicated by a star (*) mark.

## Acknowledegments

The study is supported by funds from the Chinese Ministry of Science and Technology (2017YFA0102900 ，2013CB945600), the National Natural Science Foundation of China (NSFC, 81630073 and 81372189), Science and Technology Commission of Shanghai Municipality (16JC1405700), KC Wong foundation, and Special Research Foundation of State Key Laboratory of Medical Genomics, and SJTU-USYD seed funding for Joint Research to W.Q. Gao, NSFC (81772743), the State Key Laboratory of Oncogenes and Related Genes (90-16-03), Shanghai Institutions of Higher Learning (The Program for Professor of Special Appointment (Young Eastern Scholar, QD2015002)), School of Medicine, Shanghai Jiao Tong University (Excellent Youth Scholar Initiation Grant 16XJ11003), Ren Ji Hospital (Seed Project RJZZ14-010) to H.H. Zhu.

## Author contributions

W.-Q.G. and H.H.Z conceived the study. X.W. performed most of the experiments. K.Z assisted in data interpretation. Z.Z.J. assisted in molecular cloning assays. C.P.C. and Y.R.S assited in some cell culture assays. H.F.Z assisted in confocal imaging assays. L.C.F, B.J.D, and W.X. assisted in providing tumor samples. X.W., W.-Q.G. and H.H.Z interpreted the data and wrote the manuscript.

